# Publicly Available Genomic Evidence for Nipah and Avian Influenza Viruses in Bangladesh: A Scoping Assessment of Metadata Completeness and Surveillance Representativeness at the Human–Animal Interface

**DOI:** 10.64898/2026.11.23.768755

**Authors:** Md. Nazmul Hossen, Md. Shamim Uddin, Md. Imran Hossain, Tarin Naher

## Abstract

Bangladesh experiences recurrent Nipah virus (NiV) outbreaks and sustained avian influenza virus (AIV) activity. Whether public genomic records contain metadata needed to assess surveillance representativeness across hosts, locations, and time is unclear.We conducted a structured, source-based scoping assessment of publicly documented NiV and AIV genomic records from Bangladesh (2001–2025), drawing on NCBI GenBank/INSDC and cited peer-reviewed and public-health sources. A pilot registry contained 108 entries (55 NiV, 53 AIV). Metadata were retained only when explicitly reported; unverified fields were recorded as not extracted. The registry is not a complete census. Five analytical questions were prespecified; several could not be evaluated because accession-level metadata were unavailable. The 33-accession AIV set containing H5N1 and H3N8 was excluded from H5/H9-specific interpretation. In the pilot registry, 9/108 entries (8.3%) had explicitly reported host species; the separate host-implied category was not added. Twelve entries (11.1%) had a full collection date; 21 (19.4%) had any collection date. Eight entries (7.4%) had an explicit district. The registry lists 22 NiV accession identifiers from a 2026 study, whereas the publication reports 23 newly sequenced N-gene records; this remains unresolved. Eight AIV accessions (PV961349.1– PV961356.1) represent eight segments of one chicken H5N1 isolate, not eight genomes. Twelve MN994-series records are HA-gene sequences from environmental live-bird-market samples.The pilot registry demonstrates substantial metadata limitations. Explicit host information was available for 8.3% of entries and explicit district for 7.4%; these prevent reliable quantitative assessment of host, district-level, and temporal representativeness and do not establish absence of surveillance or viral circulation. Accession-level reconstruction with transparent verification tiers and harmonised identifiers is required before quantitative representativeness indices or district-level priorities can be calculated.

## 1. Introduction

Bangladesh has experienced recurrent Nipah virus (NiV) outbreaks since 2001, with cases reported from 35 of 64 districts and a case fatality rate of approximately 71.84% [1]. The primary spillover pathway is consumption of raw date palm sap contaminated by infected Pteropus medius fruit bats, with secondary human-to-human transmission also documented [2,16]. NiV strains in Bangladesh diverge into two major lineages, NiV-BD1 and NiV-BD2, with sublineage diversification and distinct spatiotemporal clustering [3]. Phylodynamic analysis has estimated an evolutionary rate of 4.89 × 10^-4^ substitutions/site/year and documented ongoing geographic expansion, yet these inferences rest on a limited number of N-gene sequences and publicly available genomes [4].

Avian influenza viruses present a complementary challenge. Highly pathogenic AIV H5 and low-pathogenicity H9N2 are endemic in Bangladeshi poultry. A regional genomic coverage assessment found that of 919 open-access H5 isolates from nine South Asian countries, 814 (89%) originated in Bangladesh, with extreme concentration (Gini 0.83) that remained unchanged after adding restricted GISAID records [5]. H9N2 has been enzootic in Bangladeshi poultry since 2006 and exhibits complex host-species maintenance, with Bayesian phylodynamic analysis revealing frequent bidirectional viral transitions among ducks, chickens, and quails [6]. Active surveillance in live-poultry markets and the Tanguar Haor wetlands has documented reassortment of clade 2.3.4.4b A(H5N1) viruses, with the hemagglutinin gene clustering closely with LBM-origin strains and internal genes derived from contemporary duck viruses [7].

Despite this recognized burden, genomic surveillance systems remain fragmented across human and animal health sectors. A structured evaluation of country preparedness for NiV and AIV in Bangladesh identified significant gaps in detection and response capacities across sectors, and called for integrated surveillance using a One Health framework [8]. Pathogen genomics for One Health more broadly has expanded unevenly, with marked socioeconomic disparities and limited integration of multi-domain surveillance data [9]. Comparative assessments of genomic surveillance gaps for West Nile virus in Africa have demonstrated that absence of public sequences cannot be equated with absence of surveillance or viral circulation, and have called for structured frameworks that separate data availability from epidemiological risk [10]. Geospatial modelling of NiV has further shown that reservoir roost distribution and human exposure indicators extend beyond districts with recognised clinical outbreaks, underscoring the need to align genomic sampling with pathogen-specific interface indicators rather than historical case counts alone [11].

For Bangladesh, a transparent comparison of publicly documented NiV and AIV genomic records across host, spatial, temporal, and genome-completeness dimensions is needed. This study therefore has three objectives: (i) to describe the metadata available in selected public genomic records for NiV and AIV in Bangladesh; (ii) to identify where the available record structure does and does not permit comparison with epidemiological and pathogen-specific interface information; and (iii) to propose a transparent framework for future integrated genomic-surveillance assessment. A pilot registry of 108 documented accession entries is provided as a traceable starting point, not as a complete national or database-wide census.

We prespecified five analytical questions. H1 concerns the distribution of records across host and sampling-setting categories; H2 concerns the availability of district-level sequence information relative to epidemiological and ecological indicators; H3 concerns temporal continuity based on collection dates; H4 concerns sequence and genome completeness; and H5 concerns the feasibility of integrated prioritisation using these dimensions. These questions are treated as evaluative aims rather than as hypotheses assumed to be testable with the available data. Where required metadata were unavailable, the relevant question was classified as not testable or only partially assessable.

This is a scoping assessment of publicly documented evidence and a pilot accession registry, not a complete accession-level audit or a census of NCBI/INSDC and GISAID. Counts and metadata summaries below describe the assembled pilot registry only. Records reported in publications but not independently resolved at accession level are identified as lower-verification evidence. No national host proportion, district concentration statistic, temporal trend, or GSRI score is inferred from incomplete metadata.

## 2. Methods

### 2.1 Study Design and Reporting

This retrospective scoping assessment synthesised publicly documented genomic and epidemiological evidence for NiV and AIV in Bangladesh. The pilot registry was assembled from accession identifiers and accession groups reported in the cited publications and their supplementary materials. It contains 108 documented entries (55 NiV and 53 AIV), but it is not a complete database census and the verification depth differs among entries. We therefore distinguish (i) accession identifiers checked against GenBank flat files, (ii) identifiers resolved against publication supplementary tables, and (iii) records reported in publication text but not independently resolved at accession level. Quantitative statements are restricted to fields summarised in the pilot registry; unverified aggregate values from separate studies are not presented as registry-derived findings. For verification tiers, records were classified as (i) GenBank flat-file verified, (ii) resolved against publication supplementary tables, or (iii) reported in peer-reviewed text only. The three NiV records JN808857, JN808863 and JN808859 are classified as publication-text-only unless a supplement-level resolution is documented.

The study is deliberately scoped as a metadata-completeness assessment of publicly documented records; it does not perform quantitative genomic–epidemiological integration. Records that could not be resolved at accession level were retained in the descriptive inventory as unresolved rather than removed, and no accession was excluded on the basis of missing metadata alone. Accessions reported in the literature but not resolvable within the supplied sources—for example, bat-derived NiV genomes and GISAID-restricted records—are identified as not represented in the pilot registry; their absence is not interpreted as absence of surveillance or viral circulation.

### 2.2 Database Search Strategy

The assessment drew on the genomic and epidemiological sources cited in this manuscript, including NCBI GenBank/INSDC accession records where available, peer-reviewed studies, WHO Disease Outbreak News, IEDCR materials, and WOAH WAHIS reports. The supplied study documents did not preserve a complete reproducible search log containing exact database search strings, platform-specific search dates, and all screening decisions. Accordingly, this work is described as a structured source-based scoping assessment rather than a fully reproducible systematic search. The geographic scope was Bangladesh and the intended collection period was 2001–2025. The sources and accession groups included in the pilot registry are listed in Supplementary Table S3.

**Figure 1.**
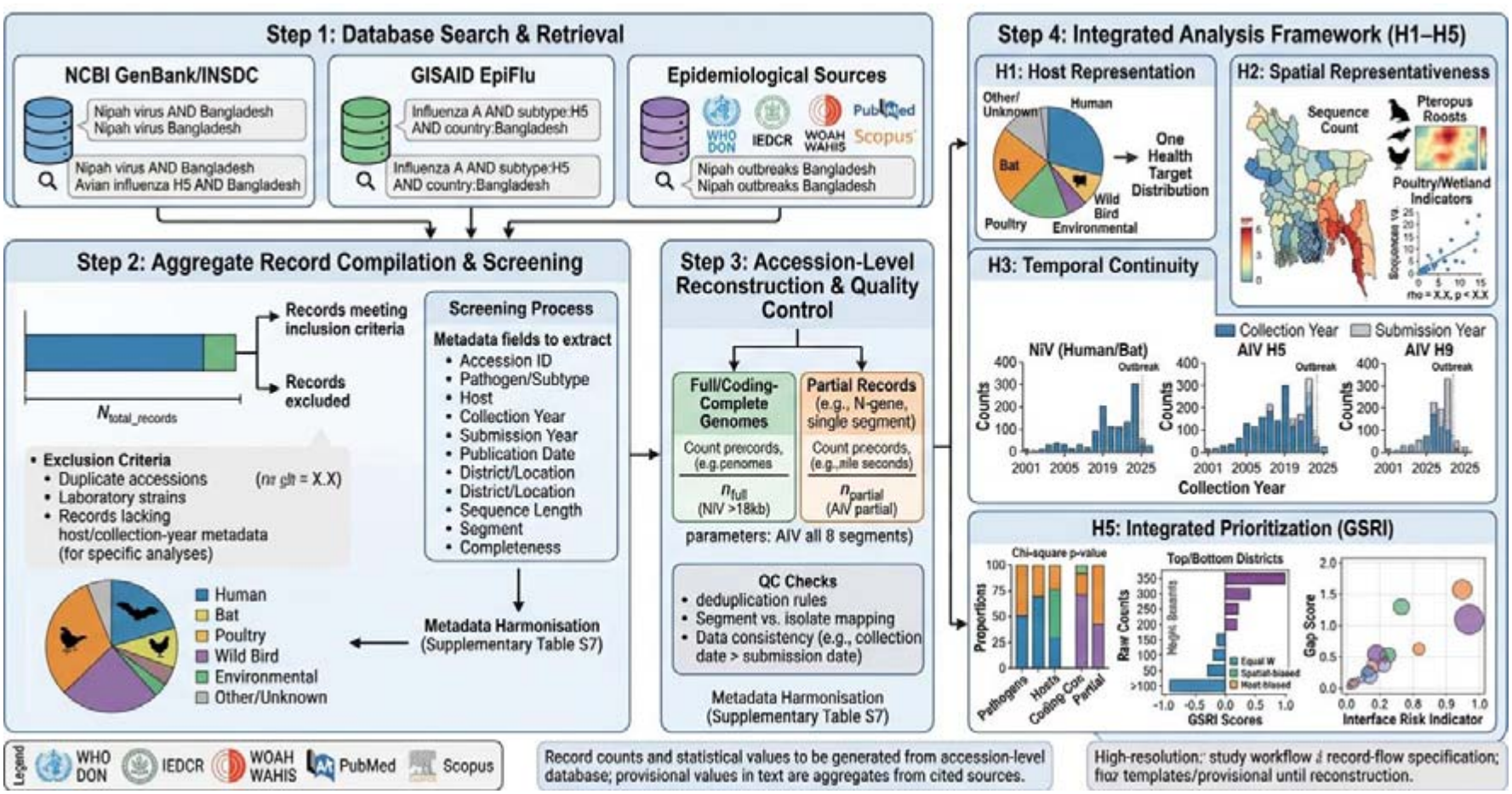
Scoping-assessment workflow and evidence-verification structure. The workflow comprises source identification, accession-list extraction, verification against GenBank flat files or publication supplements, metadata harmonisation, and eligibility assessment for pathogen- or subtype-specific interpretation. The 108-entry pilot registry is not a complete database census; counts by verification tier are reported in Supplementary Table S7. Records reported only in publication text are not labelled accession-verified.

**Table 1.** Search strategy, inclusion and exclusion criteria, and record-flow specification.

| Domain | Specification |
|---|---|
| Genomic databases | NCBI GenBank/INSDC; GISAID EpiFlu (NiV, H5, H9) |
| Epidemiological sources | WHO DON; IEDCR; WOAH WAHIS; PubMed; Scopus |
| Geographic filter | Bangladesh (country of origin) |
| Temporal filter | Collection year 2001–2025 |
| Inclusion criteria | Bangladesh-origin NiV or AIV records documented in the cited sources; accession identifier or clearly described accession group available; incomplete metadata retained for descriptive registry but excluded from analyses requiring the missing field. |
| Exclusion criteria | Exclude from a specific pathogen/subtype analysis when subtype, host, collection year, or other required variable cannot be resolved. Do not remove such entries from the descriptive pilot registry solely because metadata are missing. |
| Deduplication rule | Collapse records sharing accession ID; preserve distinct isolates and distinct genome segments; document segment-level versus isolate-level counts separately |
| Record-flow reporting | Report the 108-entry pilot registry by verification tier; distinguish accession-level verification from publication-text-only reporting. This is not a complete database screening flow. |

### 2.3 Genomic Data Extraction and Completeness

The pilot accession registry (Supplementary Table S1) contains 108 documented entries: 55 NiV and 53 AIV. Entries were included when an accession identifier or accession group was documented in the cited sources; incomplete metadata did not by itself trigger removal from this descriptive registry. Values were entered only when explicitly reported in the source material. The registry records 9/108 entries (8.3%) with an explicitly reported host species: one NiV entry (AY988601.1, human) and eight AIV accession entries (PV961349.1–PV961356.1, chicken). The separate “host implied by isolate name” category is not added to this count; if retained in the supplement, it must be reported separately and must not be merged with explicit host.The registry records 12/108 (11.1%) with a full collection date, 21/108 (19.4%) with any collection date at full-date or year level, and 8/108 (7.4%) with an explicit district. These are field-specific registry summaries and must not be interpreted as national surveillance coverage. Host and sampling setting were kept separate: live-bird-market (LBM) status denotes a sampling setting, not a bird host. The twelve MN994-series records are environmental LBM HA sequences. The eight PV961349.1–PV961356.1 accessions represent eight segments from one chicken H5N1 isolate and therefore one complete genome. The 33-accession 2023 AIV set includes both H5N1 and H3N8; because accession-level subtype mapping was not resolved, it is not used for H5/H9-specific interpretation.

Genome completeness was classified at the isolate/genome level, not at the accession-entry level. For AIV, a complete genome required all eight segments linked to the same isolate or BioSample; eight segment accessions from one isolate were counted as one genome. Records were classified as partial only when an explicit gene or segment was reported. Records reported only in publication text, or with unresolved segment/subtype mapping, were retained as unresolved and were not reclassified as complete or partial. Completeness was never inferred from isolate name alone.

Three categories of records were therefore not available for pathogen- or subtype-specific analysis: (i) the unresolved 23rd N-gene record reported by the 2026 NiV source, whose accession identifier was not available in the supplied material; (ii) the 33-accession 2023 AIV set containing both H5N1 and H3N8, for which accession-level subtype mapping was unresolved; and (iii) records reported only in publication text without accession-level verification.

### 2.4 Epidemiological and Interface Data

Human NiV records were abstracted from IEDCR surveillance summaries and WHO outbreak reports, including district, onset date, case classification, and exposure history where available [1,2]. AIV outbreak and surveillance records were abstracted from WOAH WAHIS and published active-surveillance studies covering live-poultry markets and the Tanguar Haor wetlands [7,12]. Interface indicators were pathogen-specific and prespecified. For NiV, indicators included Pteropus medius roost distribution and roost proximity to human dwellings, and documented date palm sap exposure [1,11].

For AIV, indicators included poultry density, LBM distribution, wetland coverage, and wild-bird habitat indicators [7,12]. Indicators were selected on the basis of published ecological evidence and were not combined into a single cross-pathogen score; no ecological proxy was interpreted as a direct measure of spillover probability.Epidemiological information from IEDCR, WHO Disease Outbreak News, and WOAH WAHIS was used exclusively as contextual evidence to describe the public-health and animal-health background. These data were not linked at the record level to genomic accessions. Because verified district-level sequence metadata were available for only 8/108 entries and compatible epidemiological denominators were not harmonised, no spatial join, coverage–burden correlation, or quantitative genomic–epidemiological integration was performed.

### 2.5 Statistical Framework

The following analyses were prespecified as an assessment framework. They were not all computable from the pilot registry, and no inferential statistics are claimed where the necessary record-level data were unavailable.

Host representation (H1): Host- and setting-specific proportions, confidence intervals, and diversity indices would require verified host information for each eligible accession and a clearly defined denominator. The current registry contains nine entries with explicitly reported host species. The separate host-implied category is not added to this count. Accordingly, host proportions, Shannon or Simpson diversity indices, and comparisons with a One Health target distribution were not calculated. Any equal-weight target would be a heuristic benchmark requiring biological and policy justification, not an established optimal sampling distribution.

Spatial representation (H2): District-level sequence counts, Gini coefficients, coverage–burden correlations, and spatial regression require accession-level district data and harmonised epidemiological denominators. Only eight of 108 registry entries contain an explicit district. A spatial join was not performed, and no district-level genomic concentration statistic or coverage–burden association is reported. Published outbreak counts and ecological indicators are treated as contextual evidence only and are not substituted for sequence-derived district counts. If future count modelling is undertaken, overdispersion should be assessed and the model choice justified [13].

Temporal continuity (H3): Collection date, collection year, and database submission date are distinct variables. The registry contains 12 entries with a full collection date and 21 with any collection date at full-date or year level. The available material is insufficient to reconstruct annual sequence counts or collection-to-submission lags for the full registry. Therefore, no annual trend model, zero-year analysis, or inferential temporal comparison was performed.

Genomic completeness (H4): Completeness was described only where the source explicitly identified the sequence or segment. For AIV, a complete genome requires all eight segments linked to the same isolate; eight segment accessions from one isolate count as one genome. Single-gene or single-segment records are partial genomic representations, not complete genomes. Because completeness remains unresolved for 65/108 entries (60.2%) in the pilot summary, no comparative hypothesis test across pathogens or hosts was performed. Completeness was classified at the isolate/genome unit: PV961349.1–PV961356.1 represent one complete H5N1 isolate/genome; MN994 records are HA-only partial records; the 33-accession mixed H5N1/H3N8 set remains unresolved. Unresolved records were not assigned to complete or partial categories.

Integrated prioritisation (H5): We retain the proposed Genomic Surveillance Representativeness Index (GSRI) as a conceptual framework for future use. It was not calculated, calibrated, or externally validated in this assessment. The framework must distinguish missing information from low representativeness: an unreported host or district cannot automatically be scored as poor coverage.

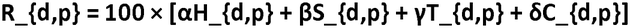

where H, S, T, and C denote host, spatial, temporal, and genomic-completeness components, respectively. Each component would require a prespecified operational definition, a valid denominator, and an explicit rule for missing or unclassifiable records.

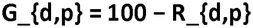

where G is a proposed evidence-gap score and higher values would indicate greater unmet evidence needs only when all component scores are estimable. Equal weights (0.25 each) and alternative weights may be explored in future sensitivity analyses, but no weighting scheme was empirically tested here. Because the current registry contains extensive missing and unresolved metadata, neither R nor G was calculated. H5 is therefore interpreted as a framework-development objective, not an empirically confirmed result.

### 2.6 Reproducibility, software, and data availability

The supplied manuscript describes R-based processing and spatial analysis as planned reproducibility procedures; however, no analysis-ready accession-level dataset or executable analysis scripts were included in the files reviewed. Software versions, package versions, search logs, and scripts should be reported only after the analyses have actually been run and archived. GISAID records remain subject to the platform’s access, attribution, and redistribution conditions and are not redistributed in this pilot registry. The GSRI was not calculated, calibrated, or externally validated. The present study does not attempt quantitative integrated prioritisation; it establishes the metadata preconditions that any such prioritisation would require.

### 2.7 Interpretation rule

Throughout, we distinguish public-data availability from surveillance activity and from pathogen circulation. The absence of an accession in the assembled registry means only that it was not resolved within the sources and extraction frame used for this pilot assessment. It does not establish that sampling, testing, sequencing, or viral circulation was absent. Ecological indicators are contextual proxies and are not interpreted as direct measures of pathogen-specific spillover probability. Epidemiological burden and interface indicators are contextual proxies only and are not statistically linked to the pilot registry. Accessions not represented in the registry include, but are not limited to, bat-derived NiV genomes and GISAID-restricted records; their absence reflects the extraction frame and platform conditions, not an absence of surveillance.

## 3. Results

### 3.1 Dataset flow and scope of provisional evidence

The pilot registry contains 108 documented entries: 55 NiV and 53 AIV. The entries were assembled from cited publications and source statements and have different levels of verification. The registry is not a complete census of public databases. Supplementary Table S7 separates GenBank flat-file verification, resolution against publication supplementary tables, and records reported only in publication text.

The results below describe what can be supported by the assembled pilot registry. They do not provide definitive estimates of national genomic-surveillance coverage. Records with unresolved subtype, host, location, collection date, or completeness are retained as unresolved rather than assigned by inference. The study is a metadata-completeness assessment, not a quantitative integration of genomic and epidemiological data. Epidemiological burden and interface indicators are reported as contextual evidence only and are not joined to the registry.

### 3.2 The pilot registry cannot support host-representation estimation (H1)

The pilot registry does not support a quantitative estimate of host representation (H1). For NiV, one complete human genome (AY988601.1) is identified in the registry and is coded as an explicit human-host entryThe 2026 source is reported to describe 23 newly sequenced N-gene records, whereas 22 accession identifiers are listed in the current registry; this discrepancy must be reconciled against the source supplement and GenBank before a final count is reported. The remaining NiV entries include records for which host or completeness is not resolved in the material assembled here. Bat-derived NiV genomes have been reported in the literature [14,15], but those records are not represented in the current pilot registry; their omission must not be interpreted as absence of bat genomic surveillance. For AIV, the 12 MN994-series entries are environmental LBM HA records, not bird-host records. Eight PV961349.1–PV961356.1 accessions correspond to the eight segments of one chicken H5N1 isolate from Gazipur in 2025 [17].The 33-accession 2023 study set contains both H5N1 and H3N8, and subtype cannot be assigned to each accession from the material reviewed; this set is therefore excluded from H5/H9-specific interpretation. With only nine entries carrying explicit host-species information, H1 is not quantitatively testable.

Several accessions and accession groups are therefore not available for pathogen- or subtype-specific interpretation, and the reasons are distinct. First, the 23rd N-gene record reported by the 2026 NiV source is unresolved because its accession identifier was not available in the supplied material. Second, bat-derived NiV genomes [14,15] were not resolved at accession level within the extraction frame and are not represented. Third, the 33-accession 2023 AIV set is excluded from H5/H9-specific interpretation because accession-level subtype mapping was unresolved. Fourth, the 12 MN994-series records are environmental LBM HA sequences and were not counted as bird-host records. Fifth, GISAID-restricted records were not redistributed and are not represented. These exclusions and non-resolutions are summarised in Supplementary Tables S1–S3 and S7.

**Table 2.**
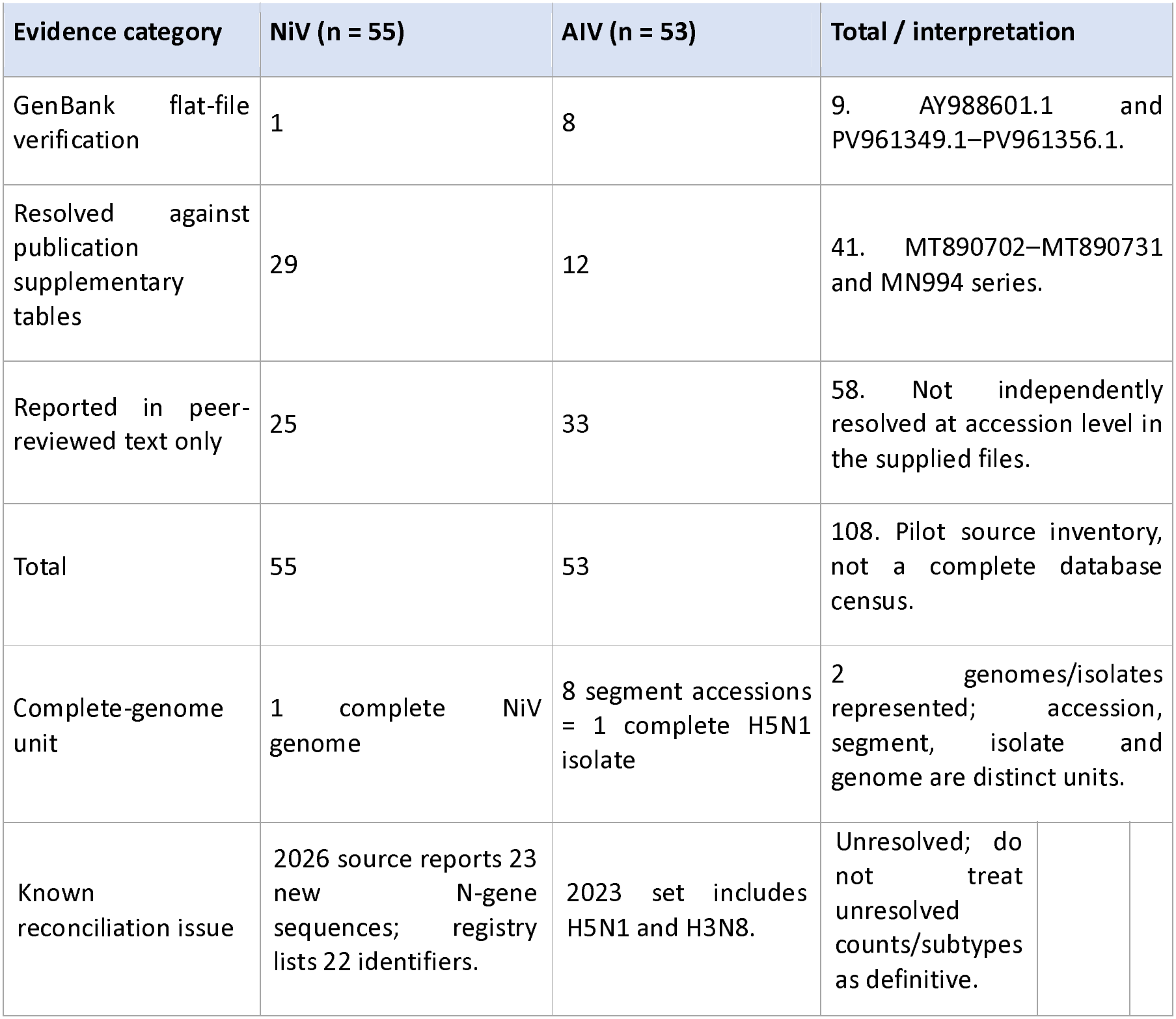
Evidence structure and verification status of the 108-entry pilot registry.

**Figure 2.**
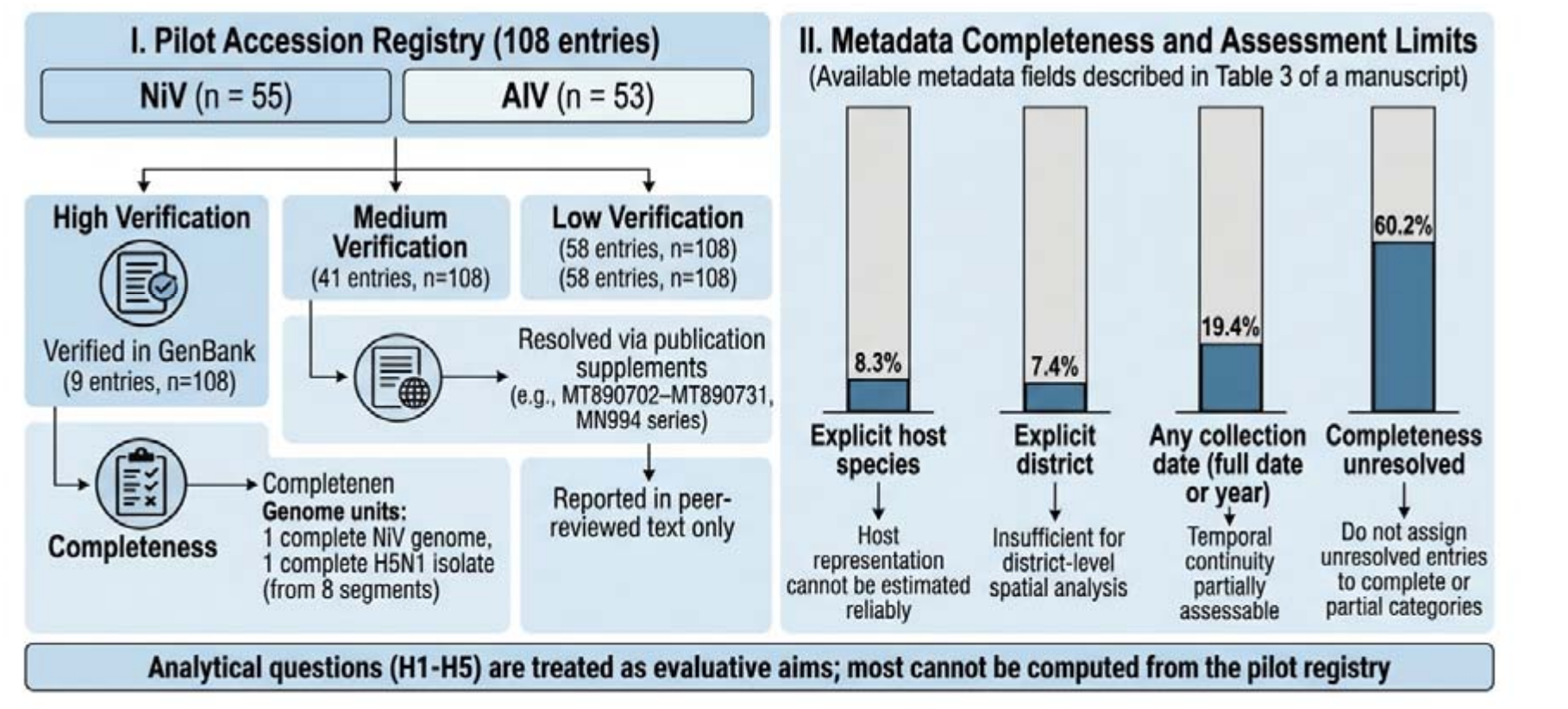
Conceptual metadata-completeness and verification-tier framework for the 108-entry pilot registry. Verification-tier counts are registry summaries from Supplementary Table S7. The figure does not display host-specific genome proportions and does not imply that the registry is a complete national census. Eight PV961349.1–PV961356.1 accessions represent one complete AIV isolate, not eight genomes.

The table summarises the pilot registry and its verification limits. Accession entries, isolates, and genome segments are different units and are not interchangeable. The 33-accession mixed H5N1/H3N8 study set is retained in the registry inventory but excluded from H5/H9-specific interpretation pending accession-level subtype resolution.

### 3.3 District metadata are too sparse to test spatial representativeness (H2)

Only 8/108 entries (7.4%) contain an explicit district in the pilot registry. Seven entries are associated with the 2016 Dhaka LBM series and one with the 2025 Gazipur chicken isolate, as recorded in the supplementary extract.District information is therefore too sparse to estimate spatial representativeness, calculate a district-level Gini coefficient, or test a relationship between sequence counts and outbreak burden. District-level outbreak counts and regional genomic concentration statistics cited from other studies are contextual findings from those sources and are not derived from this registry. H2 is consequently not testable with the current pilot data. District-level epidemiological burden and ecological indicators are presented as contextual evidence only; they were not joined to the registry and no coverage–burden association was estimated.

**Table 3.** Availability of key metadata fields in the pilot registry and limits on inference.

| Field / evidence | Registry count | Percentage | Interpretation |
|---|---|---|---|
| Explicit host species | 9/108 | 8.3% | Host representation cannot be estimated reliably. |
| Full collection date (YYYY-MM-DD) | 12/108 | 11.1% | Do not combine with year-only collection information. |
| Any collection date (full date or year) | 21/108 | 19.4% | Broader date-availability measure. |
| Explicit district | 8/108 | 7.4% | Insufficient for district-level spatial analysis. |
| Explicit division | 21/108 | 19.4% | Division information does not substitute for district coordinates. |
| Gene/segment identified | 43/108 | 39.8% | Does not by itself establish complete-genome status; includes the complete NiV genome. |
| Completeness unresolved | 65/108 | 60.2% | Do not assign unresolved entries to complete or partial categories. |

District-level outbreak and ecological information may be discussed as contextual evidence when attributed to its primary source. It must not be presented as district-level genomic coverage unless the relevant sequence accessions have verified district metadata. “No public record identified in this pilot registry” does not mean “no surveillance” or “no virus.”

**Figure 3.**
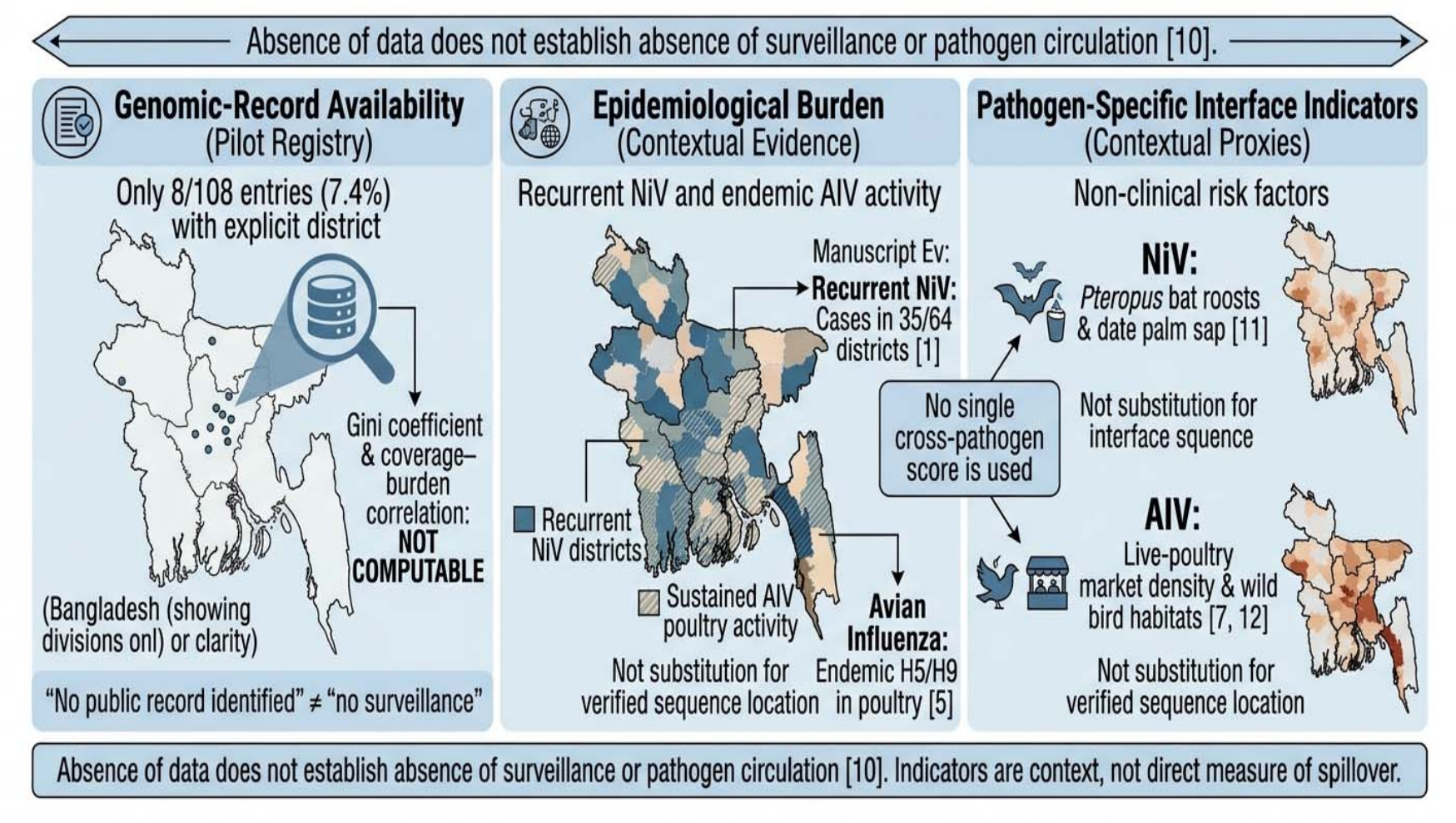
Conceptual distinction between genomic-record availability, epidemiological burden, and pathogen-specific interface indicators. Conceptual distinction between genomic-record availability, epidemiological burden, and pathogen-specific interface indicators. This framework illustration does not represent a completed district-level spatial analysis. No spatial join, Gini coefficient, coverage–burden association, or sensitivity analysis was calculated from the pilot registry.

### 3.4 Collection-date metadata are too sparse to test temporal continuity (H3)

Twelve of 108 entries (11.1%) contain a full collection date; 21/108 (19.4%) contain a collection date at either full-date or year level. These are different completeness measures and are reported separately. The current registry does not support complete annual sequence counts, collection-to-submission lag analysis, or a robust comparison with outbreak timelines. The reported gap in bat whole-genome sampling described in [14] is literature-derived context and is not directly established by the current pilot registry. H3 is therefore only partially assessable.

### 3.5 Genomic completeness is unresolved for a substantial part of the pilot registry (H4)

The registry identifies one complete NiV genome (AY988601.1) and eight AIV segment accessions (PV961349.1–PV961356.1) that collectively represent one complete H5N1 chicken isolate. The 12 MN994-series entries are HA-only environmental records. The 2023 AIV set contains 33 accessions from a study covering H5N1 and H3N8, but accession-level subtype and segment mapping were not resolved in the material reviewed. Supplementary Table S7 classifies completeness as unresolved for 65/108 entries (60.2%). Because unresolved entries cannot be safely assigned to complete or partial categories, no cross-pathogen or host-specific completeness comparison was performed. H4 is therefore descriptively assessable only for the explicitly documented subset. Completeness was classified at the isolate/genome level: one complete NiV genome and one complete AIV isolate/genome are represented; MN994 records are HA-only partial records; the 33-accession 2023 set remains unresolved. Unresolved records were retained as unresolved and were not forced into complete or partial categories.

**Figure 4.**
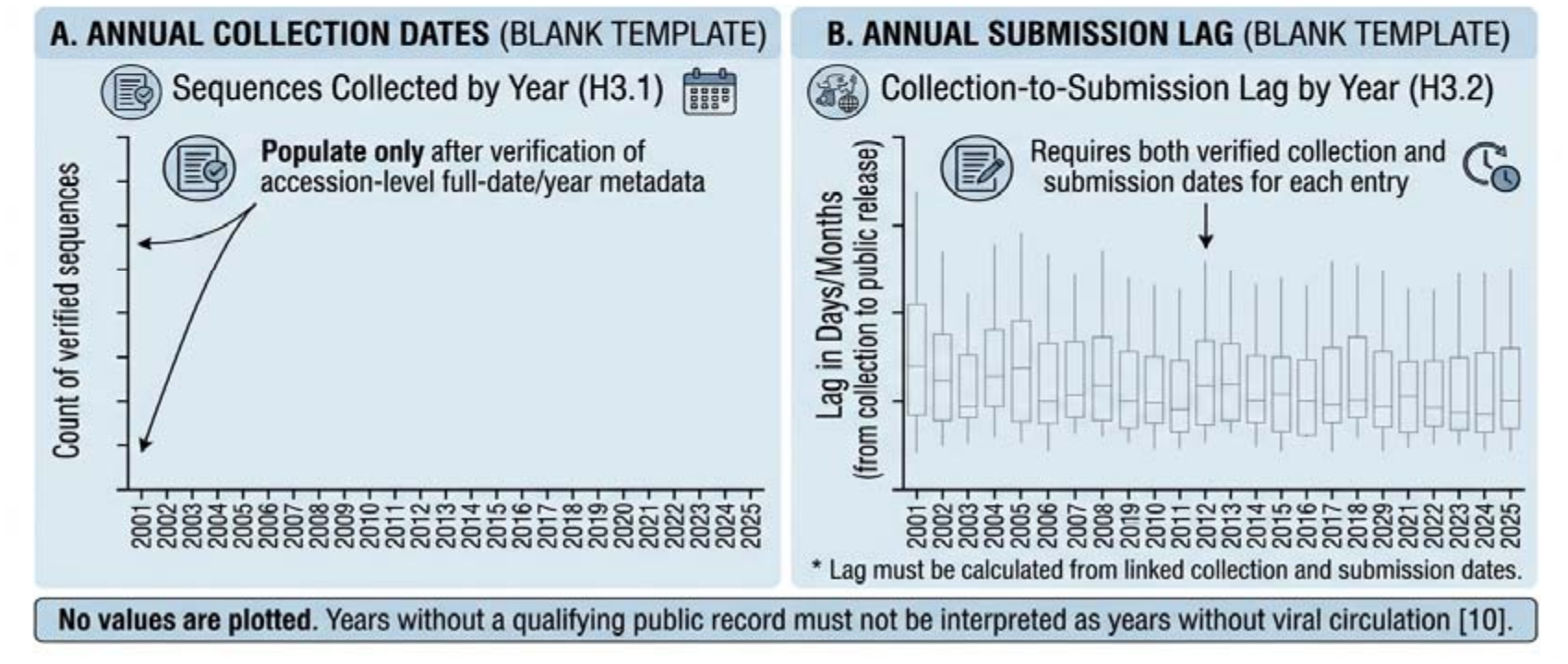
Blank templates for future annual sequence-collection and submission displays. Blank templates for future annual sequence-collection and submission displays. No values are plotted. The templates are intended to be populated only after accession-level collection and submission dates have been verified. Years without a qualifying public record must not be interpreted as years without viral circulation.

### 3.6 A minimum metadata standard is a prerequisite for integrated assessment (H5)

The current registry does not support calculation of the GSRI. The metadata summary indicates 9/108 entries (8.3%) with explicit host species, 8/108 with explicit district, 12/108 with a full collection date, and 65/108 with unresolved completeness. These are separate fields and must not be combined into a single “missing host or subtype” percentage. We therefore treat H5 as a framework-development objective: the proposed index requires verified host, location, collection date, and sequence-completeness information, as well as a defensible denominator and missing-data strategy. The GSRI remains conceptual; it was not calculated, calibrated, or validated, and no quantitative integrated prioritisation is claimed.

Proposed representativeness score: R_{d,p} = 100 × [αH + βS + γT + δC]; proposed evidence-gap score: G_{d,p} = 100 − R_{d,p}. These equations define a conceptual framework only. No score, ranking, or sensitivity analysis was calculated from the current pilot registry.

Conceptual GSRI and evidence-gap framework. The components and equations are proposed only.

**Table 4.** Proposed GSRI component definitions, direction, normalisation, and missing-data rules.

| Component | Construct | Direction | Provisional operationalisation | Target / denominator | Normalisation | Missing-data rule |
|---|---|---|---|---|---|---|
| H | Host coverage | Higher = better coverage | Host coverage relative to a justified target; not calculated in this assessment | Prespecified equal weighting across host categories | Min–max to 0–100 | Unknown host is reported separately; missingness is not scored as low coverage. |
| S | Spatial coverage | Higher = better coverage | District coverage relative to compatible burden/interface indicators; not calculated | Burden-adjusted expected coverage | Ratio to expected, capped at 100 | Districts without verified sequence location remain unassessed. |
| T | Temporal continuity | Higher = better coverage | Collection-year continuity; not calculated | Full collection period | Proportion of years covered, 0–100 | Missing collection dates are unassessed, not zero-count years. |
| C | Genomic completeness | Higher = better coverage | Proportion of complete or coding-complete genomes; not calculated | All qualifying records | Proportion, 0–100 | Unresolved completeness remains a separate category. |
| Weights | Prespecified | — | Conceptual options only; no sensitivity analysis was conducted | — | — | No ranking or composite score is reported. |

Proposed representativeness score: R\_{d,p} = 100 × [αH + βS + γT + δC]; proposed evidence-gap score: G\_{d,p} = 100 − R\_{d,p}. These equations define a conceptual framework only. No score, ranking, or sensitivity analysis was calculated from the current pilot registry.

**Figure 5.**
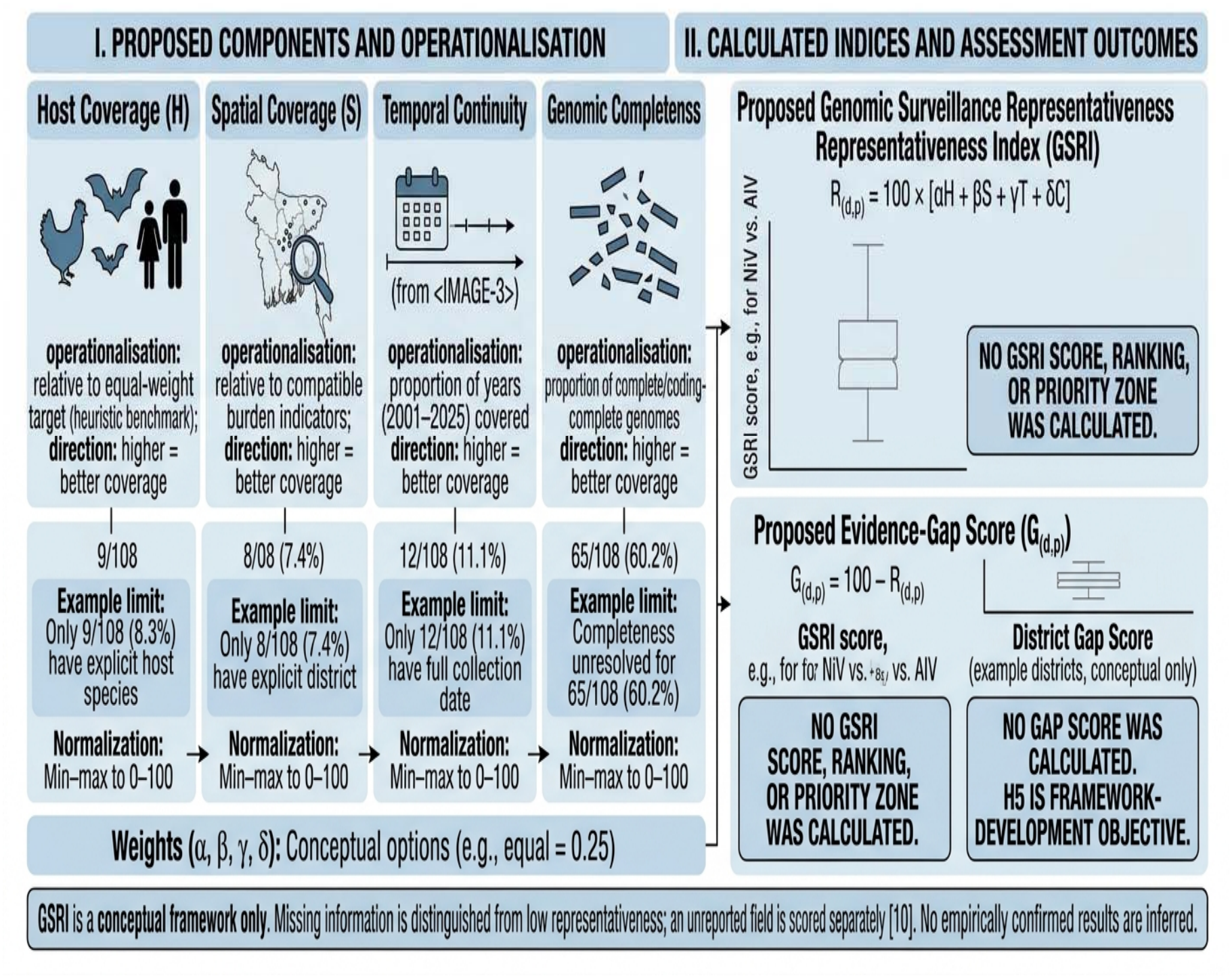
Conceptual GSRI and evidence-gap framework. No GSRI score, district ranking, priority zone, or sensitivity analysis was calculated from the pilot registry.

## Discussions

### 4.1 Principal findings

This scoping assessment identifies a central limitation in the publicly documented NiV and AIV genomic evidence assembled for Bangladesh: metadata availability is insufficient for several of the intended representativeness analyses. The pilot registry contains 108 documented entries, but it is not a complete database census and its entries have different verification levels. Explicit host, full collection-date, and district information are present in only a small fraction of entries. The current evidence therefore does not permit robust quantitative conclusions about host balance, district-level coverage, or temporal continuity. It does identify specific curation issues that must be resolved before such analyses can be undertaken.

### 4.2 Metadata incompleteness is itself the primary finding

The most defensible finding is that the pilot public-record extract is too metadata-incomplete and unevenly verified to support several planned representativeness analyses. In the 108-entry registry, 9 entries (8.3%) have an explicitly reported host species, 12 (11.1%) have a full collection date, 21 (19.4%) have any collection date at full-date or year level, and 8 (7.4%) have an explicit district. Completeness is unresolved for 65 entries (60.2%). The explicit-host count comprises one human NiV entry (AY988601.1) and eight chicken AIV segment accessions from one isolate (PV961349.1– PV961356.1). The separate implied-host category is not added to this count. Three curation issues require particular attention: (i) the 2023 AIV accession set contains both H5N1 and H3N8, so it cannot be treated as a homogeneous H5/H9 set without accession-level subtype mapping; (ii) the 12 MN994-series records are environmental LBM HA sequences and must not be coded as bird-host records; and (iii) the eight PV961349.1–PV961356.1 accessions are the eight segments of one H5N1 isolate, not eight genomes. The 2026 NiV source-reported count of 23 newly sequenced N-gene records also remains inconsistent with the 22 identifiers currently listed in the registry. These issues limit inference and should be resolved before quantitative analyses are attempted.

The distinction between accessions that are absent, unresolved, or excluded is central to interpretation. The 23rd N-gene record reported by the 2026 NiV source is unresolved because its identifier was unavailable; bat-derived NiV genomes and GISAID-restricted records are not represented because they fell outside the extraction frame or platform conditions; and the 33-accession 2023 AIV set is excluded from subtype-specific analysis because accession-level mapping was unresolved. Retaining these distinctions has direct One Health relevance: human NiV (AY988601.1), environmental LBM HA records (MN994), and a chicken H5N1 isolate (PV961) are treated as separate host, setting, and unit categories; segment counts are not conflated with genome counts; and data absence is not equated with pathogen absence. This separation supports biologically defensible interpretation across the human–animal–environment interface.

### 4.3 Biological and surveillance interpretation

For NiV, the omission of bat-derived accessions from the current pilot registry limits comparison between reservoir and human sequence evidence. Bat-derived NiV genomes have been reported in the literature [14,15], but they were not incorporated into the assembled registry. Accordingly, the current data cannot establish the extent or timing of reservoir genomic surveillance, nor can they identify where recent substitutions arose. Future work should reconcile published bat sequence reports with accessible accession records and clearly distinguish collection year, publication year, and database submission year.

For AIV, the registry illustrates why host, setting, subtype, segment, and isolate must be represented separately. Environmental samples collected in LBM settings are not equivalent to poultry-host isolates, and individual gene segments are not equivalent to complete genomes. The eight PV accessions collectively represent one H5N1 isolate, while the 33-accession study set requires accession-level subtype resolution because it includes both H5N1 and H3N8. Future surveillance should prioritise standardised metadata and transparent linkage of all eight segments to isolate or BioSample identifiers before ecological or host-specific comparisons are made.

### 4.4 Alternative explanations and interpretive caution

Several alternative explanations should be considered. Apparent gaps in the assembled public record may reflect incomplete database deposition, restricted data access, publication practices, differences in sampling effort, laboratory capacity, permitting, or metadata curation rather than an absence of surveillance or viral circulation. Historical outbreak districts may be more visible because clinical events trigger sampling, while reservoir and interface sampling may be less frequently published. Ecological convergence does not establish pathogen-specific spillover probability. No district or host should be described as pathogen-free on the basis of absent records in this pilot registry.Epidemiological burden and interface indicators are contextual and are not statistically linked to genomic records; convergence between ecological indicators and recorded outbreaks does not establish pathogen-specific spillover probability.

### 4.5 Biosafety and governance

Recommendations for field and laboratory expansion must distinguish clearly between field collection, specimen inactivation, diagnostic testing, and downstream sequencing. NiV is a high-consequence pathogen requiring appropriate containment and authorised specimen pathways. Field collection should use validated inactivation procedures before transport, and downstream processing of inactivated material should follow national and international biosafety guidance. Mobile laboratory solutions should be understood as components of a risk-based, authorised workflow—not as ordinary field facilities for handling infectious NiV material [18]. Governance arrangements should include permits, material transfer agreements, data-sharing terms consistent with GISAID attribution requirements, and community engagement at reservoir and interface sites.

### 4.6 Policy and research recommendations

We recommend five actions. First, reconcile published accession lists against GenBank records and publication supplements, documenting the verification tier and resolving the NiV 22-versus-23 discrepancy. Second, require explicit host species and sampling-setting fields, with LBM/environmental sampling kept separate from animal host identity. Third, link all AIV segments to isolate or BioSample identifiers and distinguish segment counts from isolate/genome counts. Fourth, standardise collection date, submission date, district, and division fields and publish missingness by field. Fifth, integrate verified genomic records with pathogen-specific epidemiological and ecological information only after compatible geographic and temporal denominators have been established, alongside relevant national early-warning initiatives [19,20]. Quantitative genomic–epidemiological integration should be undertaken only after verified district-level accession data and harmonised denominators are available; the present study is restricted to the metadata-completeness preconditions for such integration.

## 5. Limitations

This assessment has several limitations. It is a source-based scoping assessment and pilot registry, not a complete NCBI/INSDC or GISAID census and not a fully reconstructed accession-level dataset. The 108 entries have different verification depths: some were checked against GenBank flat files, some were resolved against publication supplementary tables, and others were reported only in publication text. The 2026 NiV source reports 23 newly sequenced N-gene records whereas 22 identifiers are listed in the registry; this remains unresolved. The 2023 AIV set includes H5N1 and H3N8, with accession-level subtype mapping unresolved, and is excluded from H5/H9-specific interpretation. The 12 MN994-series records are environmental LBM HA sequences, not bird-host records. The eight PV961349.1–PV961356.1 accessions represent one eight-segment H5N1 isolate. Explicit host, full collection date, and district information are sparse (9/108, 12/108, and 8/108, respectively), and completeness is unresolved for 65/108 entries. No district spatial join, host-diversity calculation, annual trend model, Gini coefficient, or GSRI score was calculated. GISAID-restricted records were not redistributed. Findings should therefore be interpreted as limitations of the assembled public evidence and its metadata, not as direct measures of surveillance activity, pathogen circulation, or spillover risk.

This study is a metadata-completeness assessment and not a quantitative genomic–epidemiological integration. Epidemiological data are contextual only. Genome completeness is classified at the isolate/genome level, with unresolved records retained as unresolved. The 23rd 2026 NiV N-gene record, bat-derived NiV genomes, GISAID-restricted records, and the 33-accession mixed H5N1/H3N8 set are either unresolved, not represented, or excluded from subtype-specific analysis; their status is reported rather than inferred.

## 6. Conclusion

The 108-entry pilot registry documents publicly reported NiV and AIV accessions from Bangladesh but does not constitute a complete database census. Its principal contribution is to identify metadata and curation limitations that currently prevent robust quantitative assessment of host, spatial, temporal, and genomic representativeness. Explicit host information is available for 8.3% of entries, explicit district information for 7.4%, a full collection date for 11.1%, and completeness remains unresolved for 60.2%. The NiV accession discrepancy and the mixed H5N1/H3N8 AIV study set require resolution before pathogen-specific counts are finalised. These public-record limitations do not establish absence of surveillance or viral circulation. Future analyses should begin with accession-level verification, consistent metadata standards, and clear separation of host, setting, segment, isolate, and genome units. The proposed GSRI remains conceptual and should not guide policy until it can be calculated from sufficiently complete data and externally evaluated.

The study is deliberately limited to a metadata-completeness assessment; it does not claim quantitative integration of genomic and epidemiological data. Accessions that are unresolved or not represented are reported as such, and their absence is not interpreted as absence of surveillance or viral circulation.

## List of abbreviations

AIV: avian influenza virus
GSRI: Genomic Surveillance Representativeness Index
HA: hemagglutinin
IEDCR: Institute of Epidemiology, Disease Control and Research
INSDC: International Nucleotide Sequence Database Collaboration
LBM: live-bird market
NCBI: National Center for Biotechnology Information
NiV: Nipah virus
WAHIS: World Animal Health Information System
WOAH: World Organisation for Animal Health.

## Declarations

### Author Contributions

MNH: Writing – original draft, Project administration, Writing – review & editing, Visualization, Conceptualization. MSU: Writing – review & editing, Supervision, Conceptualization. MIH: Supervision, Writing – review & editing. TN: Writing – review & editing, Project administratio

### Acknowledgment

The authors would like to acknowledge Habiganj Agricultural University and National University, Bangladesh, for institutional support and the Faculty of Veterinary Medicine and Animal Science and the Department of Paraclinical Science for promoting interdisciplinary academic collaboration. The authors also acknowledge veterinarians, public health professionals, and ecologists working in tropical settings whose field-based knowledge and broader scientific contributions have informed the context of this review.

### Ethics approval and consent to participate

Not applicable. This study used only publicly available genomic records and published literature; no human participants, human tissue, or animal experimentation were involved.

### Consent for publication

Not applicable.

## Declaration of AI-Assisted Tools

The manuscript was initially written by the authors. DeepSeek was used to assist with manuscript structure refinement, grammar correction, language polishing, and sentence restructuring. Canva was primarily used for the design, editing, and formatting of the figures, while Google Gemini was used to generate preliminary visual representations of conceptual frameworks and diagrams based on scientific concepts, information, and detailed instructions developed and provided by the authors. The Gemini-generated figures were subsequently imported into Canva for further editing and formatting. All AI-assisted text and visual outputs were critically reviewed by the authors, who take full responsibility for the accuracy, integrity, and final content of the manuscript.

## Data Availability

The pilot registry (108 documented entries) is supplied as Supplementary Table S1. It records source-derived accession identifiers and metadata, with unverified fields retained as not extracted. Supplementary Tables S2–S7 document extraction decisions, source attribution, verification tiers, and metadata completeness. The registry is not a complete NCBI/INSDC or GISAID census. GISAID-restricted records are not redistributed. Any subsequent accession-level reconstruction and analysis-ready dataset should be deposited in a public repository only after reconciliation and in accordance with database terms.

## Code Availability

No executable analysis scripts or inferential statistical outputs are reported as completed in this scoping assessment. R scripts for future accession-level curation, metadata summaries, spatial analysis, and GSRI evaluation should be deposited with the reconciled dataset when those analyses have been conducted and checked.

