## Supplementary material for "Publicly Available Genomic Evidence for Nipah and Avian Influenza Viruses in Bangladesh: A Scoping Assessment of Metadata Completeness and Surveillance Representativeness at the Human–Animal Interface": Supplimentary files

**Supplementary File**

**Table S1. Pilot accession registry: source-group inventory**

| **Registry group** | **Virus/subtype** | **Entries** | **Source-derived description** | **Verification / interpretation** |
| --- | --- | --- | --- | --- |
| 1 | NiV-B | 1 | AY988601.1; complete Bangladesh NiV genome; human host is coded as explicit; Rajbari location is mentioned in the publication narrative but is not coded as verified district metadata in the registry. | GenBank flat-file level verification reported. |
| 2–4 | NiV-B | 3 | JN808857, JN808863, JN808859; outbreak-associated records; completeness unresolved in supplied extract. | Publication text only; not independently resolved at accession level. Do not describe as supplement-resolved or as complete genomes. |
| 5–33 | NiV-B | 29 | MT890702–MT890731; accession series reported by 2021 NiV study. | Publication supplementary-table level; exact per-record metadata not fully resolved in the supplied summary. |
| 34–55 | NiV-B | 22 | PP981665–PP981684 plus PQ368170.1 and PX984254.1 as listed in the supplied registry; N-gene records. | Registry lists 22 identifiers; source is reported to describe 23 newly sequenced records. Reconciliation required. |
| 56–88 | AIV | 33 | OK/OL/ON accession set from 2023 study covering H5N1 and H3N8. | Subtype mapping not resolved accession by accession; exclude from H5/H9-specific interpretation. |
| 89–100 | AIV H5/H9 | 12 | MN994-series HA accessions from environmental live-bird-market samples in Dhaka (2016). | Environmental sampling setting; not bird-host records; HA-only. |
| 101–108 | AIV H5N1 | 8 | PV961349.1–PV961356.1: PB2, PB1, PA, HA, NP, NA, MP, NS from one chicken isolate in Gazipur (2025). | Eight segment accessions constitute one complete eight-segment genome from one isolate. |

*S1 footnote: AY988601.1 contributes one explicit-host entry (human) and one complete-genome unit. It is not added to the district-coded count because the Rajbari location is narrative-level rather than accession-level verified. JN808857, JN808863 and JN808859 are classified as publication-text-only in line with the main text and Table 2 of the manuscript.*

**Table S2. Extraction audit and required corrections**

| **Audit item** | **Finding in supplied material** | **Revised interpretation / action** |
| --- | --- | --- |
| Pilot registry size | 108 entries: 55 NiV and 53 AIV. | Retain as a pilot source inventory, not a complete database census. |
| 2026 NiV sequence count | Registry lists 22 accession identifiers; source is reported to describe 23 newly sequenced N-gene records. | Do not select either number as final until source Supplementary Table S1 and GenBank are reconciled. |
| 2023 AIV accession set | 33 accession identifiers originate from a study covering H5N1 and H3N8; accession-level mapping is unresolved. | Retain in inventory but exclude from H5/H9-specific analysis unless each accession is mapped. |
| 2016 MN994 series | 12 HA accessions are described as environmental live-bird-market samples. | Code LBM as sampling setting; do not code as chicken, duck, or other bird host. |
| 2025 PV961349.1–PV961356.1 | Eight accessions correspond to PB2, PB1, PA, HA, NP, NA, MP, and NS. | Count as eight segments, one isolate, and one complete genome. |
| Metadata completeness | Full collection date 12/108; any date 21/108; explicit district 8/108; explicit host 9/108. | Report each field separately; do not merge full-date and year-only records. |
| Verification tiers | 9 GenBank flat-file level; 41 resolved against publication supplements; 58 reported in peer-reviewed text only. | Show verification tier for each record and do not describe all 108 as independently accession-verified. |

*S2 footnote: The explicit-host count comprises one human NiV entry (AY988601.1) and eight chicken AIV accession entries (PV961349.1–PV961356.1). The separate “host implied by isolate name” category is reported separately in Table S7c and is not added to this count.*

**Table S3. Source map and verification tiers**

| **Source group** | **Virus** | **Registry count** | **Evidence described in supplied files** | **Caveat** |
| --- | --- | --- | --- | --- |
| PMC3366751 | NiV | 1 | AY988601.1; complete Bangladesh NiV genome; human host coded. | One accession. Rajbari location narrative only, not accession-level verified. |
| PMC3310473 | NiV | 3 | JN808857, JN808863, JN808859. | Publication text only; not supplement-resolved. Not all are complete genomes; metadata require accession-level confirmation. |
| IJID 2021 NiV | NiV | 29 | MT890702–MT890731 accession series. | Publication supplementary-table resolution. |
| Microbiol Spectr 2026 [ref. 4] | NiV | 22 listed | PP981665–PP981684, PQ368170.1, PX984254.1 as supplied. | Source-reported total is 23; mismatch unresolved. |
| AIV H5N1/H3N8 study (Scientific Reports 2023 equivalent) | AIV | 33 | OK/OL/ON accession set. | Contains H5N1 and H3N8; no per-accession mapping in supplied material. |
| AIV MN994 source (Virus Evolution 2020 equivalent) | AIV | 12 | MN994-series HA records. | Environmental LBM samples; HA-only. |
| Microbiol Resour Announc 2025 [ref. 17] | AIV | 8 | PV961349.1–PV961356.1. | Eight segments from one H5N1 chicken isolate. |

*S3 footnote: The “Scientific Reports 2023” and “Virus Evolution 2020” source descriptions in earlier drafts correspond to the AIV 33-accession study set and the MN994-series HA records, respectively. Where these source descriptions differ from the numbered reference list in the main manuscript, the numbered reference list takes precedence and should be checked before submission.*

**Table S4. Quality-control rules and limitations**

| **Issue** | **Required interpretation** |
| --- | --- |
| Coverage | The registry is a pilot inventory from cited sources, not a complete NCBI/INDC or GISAID census. |
| NiV 2026 count | 22 listed identifiers do not reconcile with the source-reported 23 new N-gene sequences; unresolved. |
| Metadata gaps | Do not impute host, date, district, sequence length, or completeness. Retain NE/unresolved. |
| Host-counting rule | Host is counted per accession entry; district and division per unique isolate/location record. The two units are not interchangeable. |
| AIV segment accounting | An AIV complete genome requires all eight segments linked to the same isolate/BioSample; eight accessions can represent one genome. |
| AIV scope | The 33-entry 2023 study set includes H5N1 and H3N8. Exclude from H5/H9-specific analysis until subtype mapping is completed. |
| LBM host coding | LBM is a sampling setting. Environmental LBM specimens are not automatically poultry or wild-bird hosts. |
| GISAID | Do not redistribute restricted records; comply with access and attribution requirements. |
| Spatial analysis | No spatial join was performed. Eight explicit district entries are insufficient for robust district-level inference. |
| No derived index | No host-diversity index, district Gini, annual trend model, coverage–burden association, or GSRI score was calculated. |
| Interpretation | Absence from this pilot registry is not evidence of absent surveillance, pathogen absence, or lack of viral circulation. |

**Table S5. Verified extracts retained from the supplied registry**

| **Record group** | **Accession(s)** | **Subtype** | **Host/setting** | **Year/location** | **Gene/segment** | **Interpretation** |
| --- | --- | --- | --- | --- | --- | --- |
| MN994 series | MN994085, MN994113, MN994122, MN994088, MN994124, MN994087, MN994111, MN994086, MN994265, MN994249, MN994110, MN994116 | H9N2/H5N1/H5N1–H9N2 as listed | Environmental LBM | 2016; Dhaka division | HA | 12 environmental HA records; no bird host inferred. |
| PV961 series | PV961349.1–PV961356.1 | H5N1 | Chicken isolate | 2025; Gazipur | PB2, PB1, PA, HA, NP, NA, MP, NS | Eight segments of one complete isolate/genome. |

**Table S6. Spatial-data specification and current status**

| **Field** | **Specification** | **Current status** |
| --- | --- | --- |
| Administrative boundary | Bangladesh district boundaries; source/version must be recorded when analysis is performed. | No boundary file or spatial join is documented as completed in the supplied material. |
| Coordinate reference system | EPSG:4326 (WGS 84) may be used for geographic coordinates; projected analysis requires an appropriate projected CRS. | Specification only; no coordinates were extracted. |
| Spatial unit | District (administrative level 2); division retained as broader context. | Only 8/108 entries have explicit district information. |
| Join method | Point-in-polygon join after verified coordinates are obtained. | Not performed. |
| Missing location | Retain as unassigned / not extracted; do not infer district from broad regional narrative. | No imputation. |

*S6 footnote: The eight district-coded entries comprise seven MN994-series records from the 2016 Dhaka LBM series and one PV961 isolate from Gazipur (2025). Because PV961 is counted as one isolate for district purposes, the eight district-coded entries do not correspond one-to-one to the eight PV961 accessions.*

**Table S7. Record flow, verification tiers, subtype scope, and metadata completeness**

**S7a. Registry stage and verification tier**

| **Registry stage** | **NiV** | **AIV** | **Total** | **Interpretation** |
| --- | --- | --- | --- | --- |
| Entries in pilot registry | 55 | 53 | 108 | Source-assembled inventory. |
| GenBank flat-file verification | 1 | 8 | 9 | AY988601.1 and PV961349.1–PV961356.1. |
| Resolved against publication supplementary tables | 29 | 12 | 41 | MT890 series and MN994 series. |
| Reported in peer-reviewed text only | 25 | 33 | 58 | Not independently resolved at accession level in the supplied files. |

**S7b. AIV category scope**

| **AIV category** | **Accession count** | **Interpretation** |
| --- | --- | --- |
| H5N1 confirmed, HA-only | 2 | MN994265 and MN994249. |
| H9N2 confirmed, HA-only | 8 | MN994085, MN994113, MN994122, MN994088, MN994124, MN994087, MN994111, MN994086. |
| H5N1/H9N2 mixed reported | 2 | MN994110 and MN994116; subtype coding should remain as reported. |
| H5N1 complete eight-segment isolate | 8 accessions / 1 isolate | PV961349.1–PV961356.1; one chicken isolate. |
| Unresolved H5N1/H3N8 study set | 33 | Not eligible for H5/H9-specific interpretation until accession-level subtype mapping. |

**S7c. Metadata completeness (field-specific, not merged)**

| **Metadata field** | **NiV** | **AIV** | **Total** | **Completeness / interpretation** |
| --- | --- | --- | --- | --- |
| Explicit host species | 1 | 8 | 9/108 (8.3%) | AY988601.1 (human) and PV961349.1–PV961356.1 (chicken). |
| Host implied by isolate name | 0 | 0 | 0/108 (0%) | Not used in the current registry; reported here for completeness only. |
| Full collection date (YYYY-MM-DD) | 0 | 12 | 12/108 (11.1%) | Full date only. |
| Any collection date (full date or year) | 1 | 20 | 21/108 (19.4%) | Broader measure; do not conflate with full date. |
| Explicit district | 0 | 8 | 8/108 (7.4%) | Insufficient for district-level analysis. |
| Explicit division | 1 | 20 | 21/108 (19.4%) | Division is not a substitute for district. |
| Gene/segment identified | 23 | 20 | 43/108 (39.8%) | Does not establish complete-genome status; includes the complete NiV genome. |
| Complete-genome status | 1 | 8 segment accessions | 9 accessions; 2 genomes/isolates represented | One NiV genome and one AIV eight-segment isolate, subject to the stated verification scope. |
| Completeness unresolved | 32 | 33 | 65/108 (60.2%) | Do not reclassify unresolved records as complete or partial. |

*S7c footnotes:
1. Counting units: Host is counted per accession entry; district and division are counted per unique isolate or location record. The eight PV961 accessions contribute eight host-coded entries but only one district-coded isolate (Gazipur). The twelve MN994 accessions contribute twelve sampling-setting entries, of which seven carry explicit Dhaka district metadata.
2. Category overlap: The gene/segment-identified and complete-genome categories are not mutually exclusive. The complete NiV genome (AY988601.1) is included among the 43 gene/segment-identified entries. The sum 1 (complete) + 23 (gene/segment identified) + 32 (unresolved) = 56 therefore exceeds the NiV total of 55 because of this overlap.
3. AY988601.1: AY988601.1 is coded as an explicit human-host entry (NiV 1). Its Rajbari location is narrative-level only and is not coded as verified district metadata, consistent with the main text. If future accession-level verification confirms district, the explicit-district count should be updated to 9/108.
4. Explicit host count: The explicit-host count comprises one human NiV entry and eight chicken AIV segment accessions from one isolate. The “host implied by isolate name” row is retained here as 0/108 because no entry in the current registry is supported by isolate-name evidence alone; any future addition must be reported separately and not merged with explicit host.*
